# Extended culture of 2D gastruloids to model human mesoderm development

**DOI:** 10.1101/2024.03.21.585753

**Authors:** Bohan Chen, Hina Khan, Zhiyuan Yu, LiAng Yao, Emily Freeburne, Kyoung Jo, Craig Johnson, Idse Heemskerk

## Abstract

Micropatterned human pluripotent stem cells (hPSCs) treated with BMP4 (2D gastruloids) are among the most widely used stem cell models for human gastrulation. Due to its simplicity and reproducibility, this system is ideal for high throughput quantitative studies of tissue patterning and has led to many insights into the mechanisms of mammalian gastrulation. However, 2D gastruloids have only been studied up to 48h. Here we extended this system to 96h. We discovered a phase of highly reproducible morphogenesis during which directed migration from the primitive streak-like region gives rise to a mesodermal layer beneath an epiblast-like layer. Multiple types of mesoderm arise with striking spatial organization including lateral mesoderm-like cells on the colony border and paraxial mesoderm-like further inside the colony. Single cell transcriptomics showed strong similarity of these cells to mesoderm in human and non-human primate embryos. However, our data suggest that the annotation of the reference human embryo may need to be revised. This illustrates that extended culture of 2D gastruloids provides a powerful model for human mesoderm differentiation and morphogenesis.

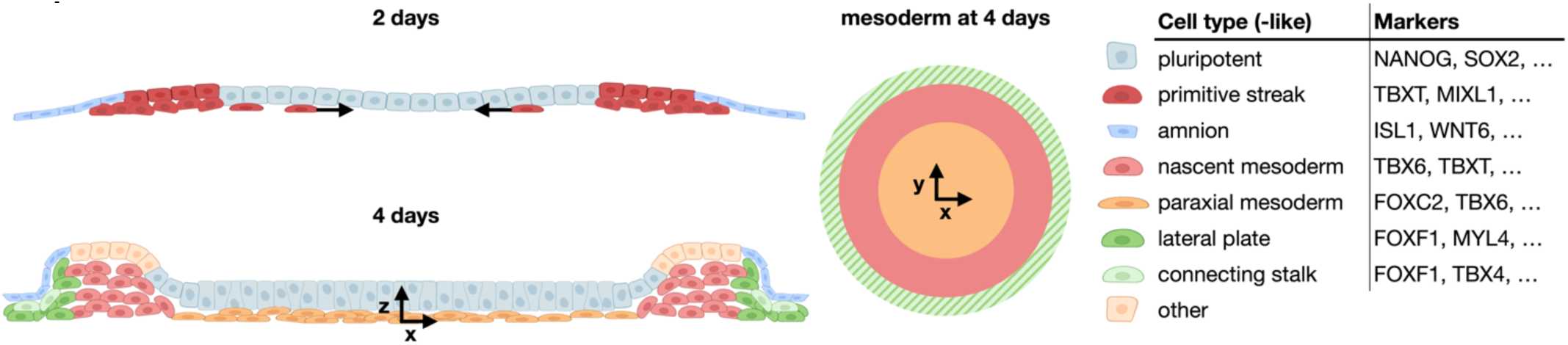
Graphical Abstract.

## Introduction

The mesoderm is one of the three embryonic germ layers that forms during gastrulation by migration from the primitive streak and is essential for the establishment of the body plan^1^. Defects in mesoderm migration may underlie some of the most common congenital birth defects, most prominently those of the heart^2,3^. Yet, despite its importance, very little is known about human mesoderm morphogenesis. Mesoderm differentiation also remains insufficiently understood. Due to the dynamic nature of the primitive streak, it remains unclear if committed subpopulations of the streak give rise to the different mesodermal lineages and if so when these lineages segregate. Addressing this is essential not only to provide insight into the etiology of human birth defects but also for stem cell technologies that require reliable differentiation to specific mesodermal lineages^4^. Recent progress in modeling aspects of human embryonic development with stem cells offers hope of investigating human mesoderm morphogenesis in vitro. However, while the latest generation of stem cell models undergo advanced morphogenesis, they display a high degree of heterogeneity and the protocols to generate them are not yet compatible with high resolution live cell microscopy and mechanical manipulation methods required to study their morphogenesis^5–7^. Therefore, simplified models are needed that capture essential behavior while at the same time enabling precise experimental control and high-resolution live cell microscopy.

2D gastruloids were among the first stem cell models for human gastrulation to be established^8^ and remain unsurpassed in reproducibility and ease of performing high resolution live cell microscopy or experimental manipulations. Consequently, they have enabled many new insights into the mechanisms of embryonic patterning^9–14^. By 42h they consist of at least four cell types, arranged concentrically: a central region of pluripotent cells surrounded by a ring of primitive streak-like cells (PSLC), then a ring of primordial germ cell-like cells, and finally an outer ring of amnion-like cells (AMLC)^8,13^. However, it has not been reported if and how this model continues to develop at later times. Therefore, we extended the culture of 2D human gastruloids from 48 to 96h. We found that highly reproducible mesoderm morphogenesis and patterning occur during this time frame, thereby establishing a powerful approach to reveal the mechanisms of human mesoderm development.

## Results

### 2D gastruloids organize into a layered 3D structure after 4 days

We discovered that 2D gastruloids remain highly organized through 4 days of culture if sufficiently high initial cell density is ensured by 24h of culture on micropatterns before BMP4 treatment (see methods). In the plane of the gastruloid (xy), the “canonical” markers at 48h: ISL1 for amnion-like cells, BRA for primitive streak, SOX2 for epiblast, were still present at 72h and 96h with roughly similar organization: ISL1+ cells on the colony edge surrounded a ring of BRA+ cells, while SOX2+ cells were present in the center (Fig.1a). However, at later times, a significant number of BRA+ cells was also observed scattered away from the ring in the colony center, most prominently at 72h (Fig.1ab, white arrows). In contrast to the relatively static appearance of the canonical markers in the xy-plane, pronounced morphological changes occurred over time in the orthogonal z-direction. At 72h, colony thickness was relatively uniform, but by 96h the edges were raised relative to the colony center, and the colony expanded slightly in the radial direction (Fig.1b, Supplementary Fig.1a). Moreover, in the colony center, a layer of cells negative for ISL1, SOX2, and BRA was consistently present underneath multiple layers of SOX2 positive nuclei by 96h. This multilayered organization with raised edges at 96h was apparent in brightfield images as a darker center and still darker ring around the edge (Supplementary Fig.1b). We developed a custom image analysis pipeline with which we were able to accurately segment individual nuclei in 3D and obtain their positions and fluorescence intensities (Methods, Supplementary Fig.1c). Single cell quantification showed that colony morphology and marker expression within the same experiment were highly reproducible (Fig.1c, Supplementary Fig.1d), although the relative height of the edges and center as well as the colony diameter varied somewhat from experiment to experiment. We confirmed this result in three different cell lines (SI Fig.1e).

The cells in the central upper layer maintained a pluripotent, epiblast-like identity since SOX2 positive cells co-expressed NANOG and OCT4 (Fig.1e). Staining for the epithelial markers EPCAM and ECAD and the mesenchymal markers SNAI1, NCAD, and VIM, revealed continuous epithelium covering the top of the colony with mesenchymal cells underneath (Fig.1fg). An additional mesenchymal population was present outside the streak-like region on the colony edges (Fig.1fg). The epithelial apical surface marker PODXL was present only on the colony surface indicating that the multiple layers of pluripotent nuclei in the center belong to a single pseudo-stratified epiblast-like layer with mesenchyme underneath (Fig.1g).

The scattered appearance of BRA+ cells in the center at 72h suggests that the mesenchyme is mesoderm that forms by cells migrating out of the primitive streak (PS)-like ring and subsequently downregulating PS markers. We therefore investigated PS marker expression more closely and discovered unexpected heterogeneity. Although BRA, EOMES, and MIXL1 stains appear very similar (Supplementary Fig.1fg), their overlay and quantification revealed graded expression with BRA higher outside and EOMES and MIXL1 higher inside at 48h and 72h (Fig. 1hi). This is consistent with their distinct roles in posterior and anterior primitive streak, respectively, as BMP signaling is higher on the colony edge which thereby resembles the posterior of the embryo^9,15,16^. Heterogeneity at the single cell level increased over time and different PS markers lost the strong correlation they exhibit at 48h (Fig. 1jkl). In particular, anterior and posterior streak markers separated further and BRA+MIXL1-EOMES– and BRA-MIXL1+EOMES+ populations were present at 96h. Although in the streak-like ring these populations had lost the spatial organization present at 48h and appeared randomly mixed, the cells in the colony center belonged exclusively to the BRA-MIXL1+EOMES+ population. This suggests that distinct PSLC subpopulations may give rise to a central mesodermal layer and mesodermal ring on the colony edge.

### Distinct mesodermal populations arise by 96h

To determine if the observed mesenchymal populations indeed represent distinct mesodermal cell types, we performed single cell RNA-sequencing at 48, 72, and 96h in duplicate. We created an integrated UMAP visualization and clustered the data using the Leiden algorithm with parameters chosen to produce clusters at 48h that were consistent with those previously obtained and verified at 42h^13^ (Fig. 2ab). In agreement with Fig.1h-l this yielded distinct primitive streak-like clusters, respectively marked by co-expression of TBXT, MIXL1, and EOMES and another by TBXT and WNT8A in the absence of MIXL1 and EOMES (Fig. 2bc). We tentatively named these anterior and posterior streak-like cells (APS-LC and PPS-LC). In addition, we found two major mesodermal clusters.

Differential expression analysis yielded markers including TBX6, DLL3, RSPO3, FOXC1 for one cluster and FOXF1, HAND1, TMEM88 for the other (Fig. 2c, Supp. Data 1). In the CS7 human gastrula^17^, similar signatures were associated with clusters respectively annotated Nascent and Advanced Mesoderm, and we named them accordingly. Nascent mesoderm was already present at 48h, consistent with^18^, but grows to the largest population by 96h, while advanced mesoderm was first seen at 72h and mostly specified between 72 and 96h (Supp.Fig.2.a). We then clustered the 96h data separately, which further subdivided the mesoderm (Fig. 2d). The nascent mesoderm was split into what appeared to be an early and late cluster, which, in keeping with the literature we could have named nascent and emergent mesoderm. However, the later cluster showed downregulation of MIXL1 and RSPO3 while a fraction of cells highly expressed FOXC2 and MIXL1, consistent with the transition from early to late paraxial mesoderm^19,20^ (Fig.2e). We therefore named the later cluster paraxial mesoderm-like cells (PM-LC). The advanced mesoderm separated into a cluster expressing markers of lateral plate– and early cardiac mesoderm--which we therefore named LPM-like cells (LPM-LC)–– and a cluster marked by TBX4, PITX1, MLLT3 and ALCAM (Fig. 2e). In the mouse gastrula, TBX4 is exclusive to the allantoic mesoderm^21^. Moreover, the yolk-sac mesoderm markers POSTN, DCN and ANXA1 were not (or minimally) expressed. We therefore hypothesized that this cluster represents the extraembryonic mesoderm of the connecting stalk (CS-LC). In the embryo this tissue is in contact with the amnion, consistent with the fact that amnion is present in 2D gastruloids, enabling tissue interactions that may give rise to CS mesoderm^22,23^.

To verify the identity of our cells, we then mapped the gastruloid transcriptome onto the CS7 human gastrula^17^ (Fig 2fg). As expected, our advanced mesoderm cluster mapped to the CS7 advanced mesoderm, while nascent mesoderm mapped predominantly to nascent mesoderm with small discrepancies likely due to somewhat arbitrary boundaries between discrete clusters on a transcriptional continuum (Fig. 2h-j). Since both our LPM-like and CS-like cells mapped to the advanced mesoderm, we verified these identities by also comparing to the CS8 cynomolgus monkey embryo, in which LPM, allantois, and (somatic) extraembryonic mesoderm (ExMe) were separately annotated^24^ (Fig.2k). Consistently, our LPM-LCs mapped to the LPM, while the CS-LCs mapped around the borders between the clusters annotated as ExMe, allantois, and pharyngeal mesoderm (Fig.2lm, Supp.Fig.2b). Marker gene analysis confirmed that this region matches the gene expression profiles of our CS-LCs: TBX4+PITX1+ISL1+ANXA1-DCN-(Supp.Fig.2c-e).

Our finding suggests that the cells annotated as advanced mesoderm in the CS7 human embryo may also be a mixture of lateral plate and connecting stalk mesoderm. Consistent with this, the sample contained the connecting stalk and somatic extraembryonic mesoderm (Tyser^17^ Fig.1) but these cell types were not annotated in the scRNA-seq data. Moreover, like in the gastruloids, the advanced mesoderm from the human gastrula contained mutually exclusive regions expressing PITX1 on one side and MYL7, MYL4 on the other (Supp.Fig.2d). Although TBX4 expression was lacking, this may be due to sequencing depth or developmental stage, as it not clear when this marker is first expressed. Mapping the human CS7 data to the monkey CS8 and transferring the monkey annotation back to human further supports the conclusion that the advanced mesoderm is composed of lateral and somatic extraembryonic mesoderm (Fig.2n).

**Figure 1:**
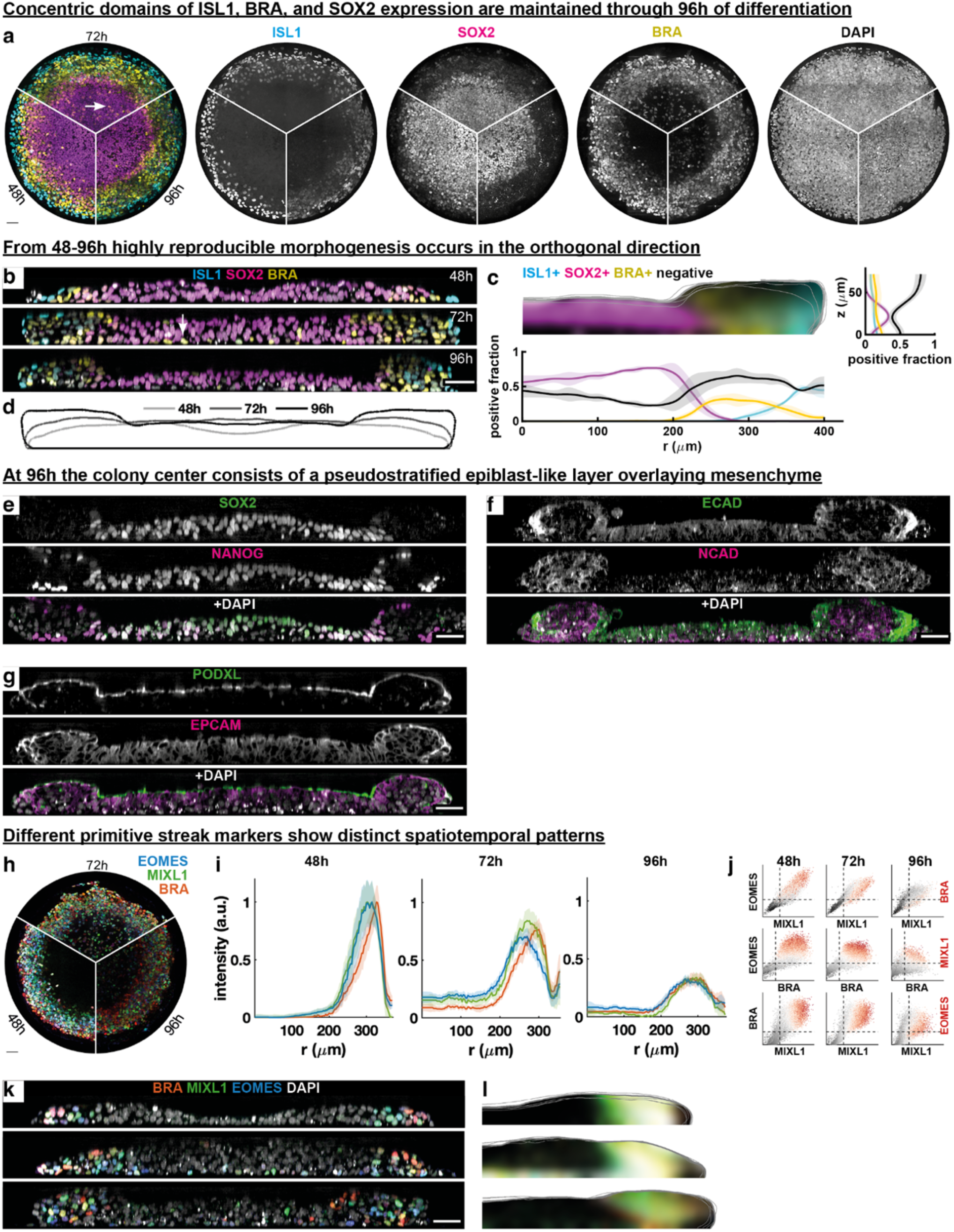
2D gastruloids organize into a layered 3D structure after 4 days. (**a**) Maximal intensity z-projection (MIP) of 2D gastruloids at 48, 72, and 96h after BMP4 treatment. **(b)** Cross-sections through colony centers in (a). Arrows in (a,b): BRA+ cells in colony center. **(c)** Combined distribution for N=4 colonies of cells positive for indicated markers at 96h as a function of r and z. **(d)** Average outline at different times for N=4 colonies per time. **(e-g)** Cross-section of stainings for pluripotency markers (e), apical marker PODXL and epithelial marker EPCAM (f), E-cadherin and N-cadherin g). **(h)** MIP of immunofluorescence stain for 3 primitive streak (PS) markers. **(i)** Radial intensity profiles of primitive streak markers at different times. **(j)** Scatterplots of primitive markers over time. **(k)** Cross-sections corresponding to (h). Note that PS-like cells in the center at 72h are high MIXL1 or EOMES, low BRA, consistent with (i) showing low BRA in the center at 72h. (**l)** Distributions of positive cells over N=4 colonies.

**Figure 2:**
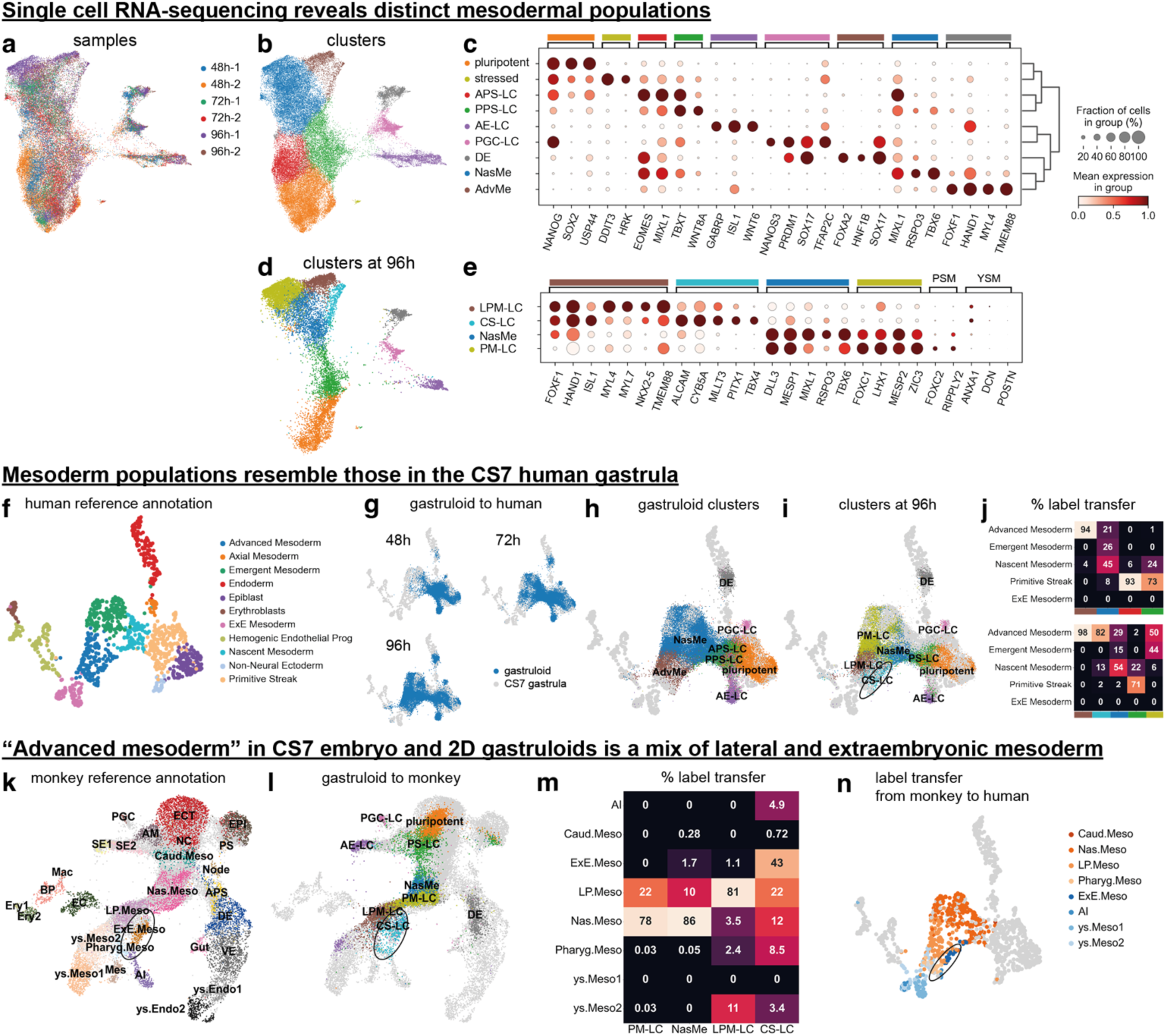
Distinct mesodermal populations arise by 96h. (**a**) Integrated UMAP for gastruloid time series single cell RNA-sequencing. **(b)** Annotated Leiden clusters. –LC: like-cells, APS: anterior primitive streak, PPS: posterior primitive streak, AE: amniotic ectoderm, PGC: primordial germ cell, DE: definitive endoderm, NasMe: nascent mesoderm, AdvMe: advanced mesoderm. **(c)** Dot plot of normalized marker genes expression in different clusters. **(d)** Leiden clusters for 96h samples. LPM: lateral plate mesoderm, CS: connecting stalk, PM: paraxial mesoderm. **(e)** Dot plot of normalized marker gene expression in mesodermal clusters at 96h. **(f)** Annotated clusters for CS7 human gastrula^17^. **(g)** Gastruloids at different times mapped onto reference CS7 gastrula. **(h)** Gastruloid clusters mapped on to gastrula. **(i)** Gastruloid clusters at 96h mapped onto gastrula, circle shows CS-LC and LPM-LC mapping to separate regions of the gastrula advanced mesoderm. **(j)** Percentage of cells from each gastruloid mesodermal cluster mapped to gastrula mesodermal clusters using KNN label transfer. **(k)** Annotated clusters for CS8 monkey embryo^24^. **(l)** Gastruloid clusters at 96h mapped onto monkey embryo. Circle indicates mapping of CS-LCs. **(m)** Percentage of cells from each gastruloid mesodermal cluster mapped to monkey mesodermal clusters using KNN label transfer. **(n)** KNN label transfer from mesodermal clusters to human gastrula shown on gastrula UMAP with embryonic in shades of red and extraembryonic in shades of blue. Circle shows extraembryonic mesoderm mapping to the same region of the advanced mesoderm as the gastruloid CS-LCs.

### Mesodermal populations are organized from edge to center by directed migration

Immunofluorescence staining confirmed the existence of distinct cell populations marked by TBX6 and FOXF1 and revealed a clear spatial organization. TBX6+ mesoderm formed a layer underneath the epiblast-like cells in the center (Fig.3a-c, Supp.Fig.3a). In addition, TBX6+ cells formed a ring underneath the raised part of the colony in the position of the streak-like ring at 48h (Fig.3a), consistent with the expression of TBX6 in the mid to late primitive streak^25,26^. In contrast, FOXF1 positive mesoderm formed on the colony edge (Fig.3d-f, Supp.Fig.3b). Consistent with the scRNA-seq, the HAND1 expression domain overlapped with both FOXF1 and TBX6 (Fig.3g-i, Supp. Fig.3c). This spatial organization of mesodermal populations was reproducible across multiple cell lines (Supp. Fig.3d-i).

We next determined the spatial organization of mesodermal subpopulations. We discovered that the mesodermal layer underneath the epiblast co-expresses FOXC2 with TBX6, while the peripheral TBX6+ ring does not express FOXC2, suggesting that the ring may represent the late streak or nascent mesoderm, while the cells that migrate out of the streak to the center are differentiated paraxial mesoderm (Fig.3j-l, Supp.Fig.3j). We also confirmed that subsets of the FOXF1+ population expressed the early cardiac marker NKX2.5 (Fig.3mn, Supp.Fig.3k) and CS marker ALCAM (Fig.3op, Supp.Fig.3l), with NKX2.5 more on the bottom edge and ALCAM further inward and more on top, close to the ISL1+ amnion. Altogether, these data show that the mesenchymal layer in the center and along the colony edge represent distinct mesodermal populations.

The observation of primitive streak-like cells outside the streak at 72h suggests that the mesodermal populations originate in the streak-like region, from which they migrate to the colony center and edge. Consistent with this, live cell imaging showed BRA+ cells migrating out of the streak-like ring starting around 48h, although the onset of migration varies somewhat from experiment to experiment (Supp. Video 1). Furthermore, single cell tracking of BRA-positive cells showed that nearly all BRA+ cells in the center at 61h originate from the primitive streak-like ring at 36h (Fig. 3qr, Supp.Fig.4a). Moreover, the inward migration of the BRA+ cells in the center is directed, as the probability of cells to move radially inward is much greater than any other directions (Fig. 3s). In addition to inward migration that forms the mesodermal layer in the center, we observed outward migration toward the colony edge, consistent with the idea that the FOXF1+ population on the colony edge also originates from the streak (Supp.Fig.4b-c). In contrast to the inward migration where cells appear to mostly migrate individually, outward migration may be collective (with groups of cells migrating together), followed by tangential migration along the colony edge (Supp.Fig.4d). In summary, our data suggests that similar to the embryo, different mesodermal populations in 2D gastruloids migrate from the streak-like region in a directed and spatially organized manner that resembles movements seen in the embryo.

**Figure 3:**
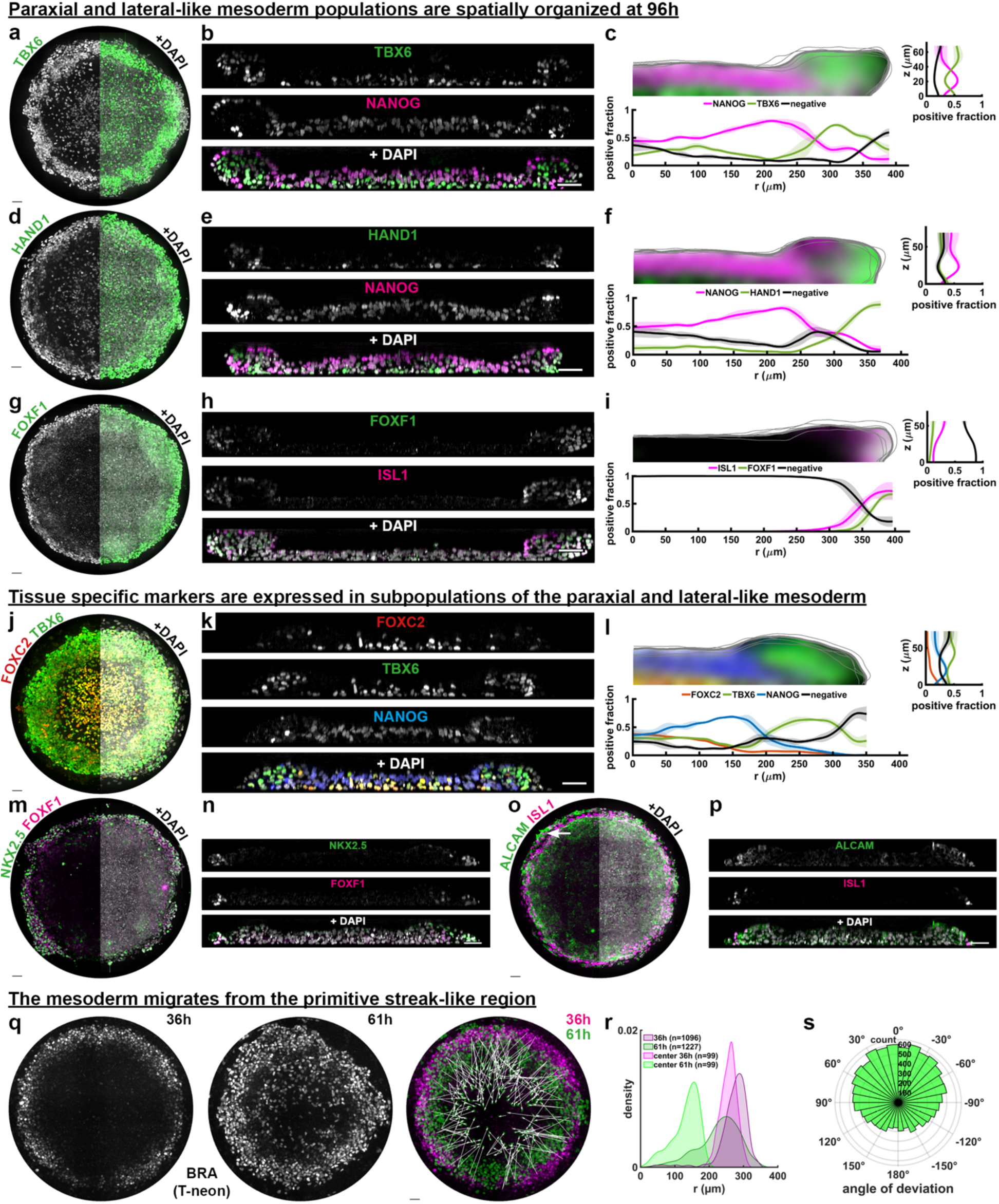
Mesodermal populations are organized from edge to center by directed migration. (**a-i**) Immunofluorescence stainings for TBX6, FOXF1 and HAND1 at 96h shown as **(a,d,g)** maximal intensity projection along z without (left) and with DAPI (right), **(b,e,h)** cross sections through the colony, and **(c,f,i)** quantification of the average fraction distribution of cells positive for each marker as a function of r and z. **(j-p)** Like (a-i), for paraxial (FOXC2), early cardiac (NKX2.5), connecting stalk-like (ALCAM+) sub-populations. Arrow in (o): high ALCAM+ cells within the lateral-like mesoderm. **(q)** H9 T-neon (BRA) colony at 36h (left) and 61h (also see Supplementary Video 1) and overlay of 36h and 61h with white arrows showing the total displacement of tracked cells whose final positions are in the colony center. **(r)** Histogram of the initial and final radial distribution of all BRA+ cells and BRA+ cells that are in the center in the final frame (see methods). **(s)** Histogram of the angle of deviation from radially inward motion for all track segments (Supp.Fig.4a) for cells tracked to the center in (o). Scale bars 50um. Source data are provided in a Source Data file.

### Mesodermal populations depend on distinct signaling activity

Lateral mesoderm depends on high BMP signaling, while paraxial mesoderm depends on Wnt signaling^19^. This suggests the spatial patterning of the mesoderm may depend on graded BMP signaling from edge to center. To determine BMP signaling activity at 96h we stained for pSMAD1, which was indeed higher on the outside of the colony including the FOXF1+ domain (Fig.4a, Supp.Fig.5a,c). However, the domain of high pSMAD1 extended well past the FOXF1+ domain and covered the entire raised colony edge. To test the role of BMP signaling we removed BMP from the media after 48h, which only had a minimal effect (Fig.4a, Supp.Fig.5a,c). The size of the raised edge and pSMAD1 levels in the colony center were slightly reduced, but BMP signaling remained high in the edge region, suggesting that the primary source of BMP ligands at this stage is endogenous. Consistently, the mesoderm populations express high levels of BMP2 and BMP4 (Supp.Fig.5d). To eliminate BMP signaling we then treated the colonies with a BMP receptor inhibitor, which nearly eliminated FOXF1 expression (Fig.4a, Supplemental Fig.5a,c) In contrast, removal of BMP and inhibition of BMP signaling did not reduce TBX6 expression and instead led to a slight outward expansion of the TBX6+ population (Fig. 4b, Supp.Fig.5b,c). We also observed a significant change in NANOG expression with BMP receptor inhibition. This was due to a loss of primordial germ cell-like cells under these conditions, while NANOG expression in the pluripotent epithelium was unaffected (Supp.Fig.5de).

**Figure 4:**
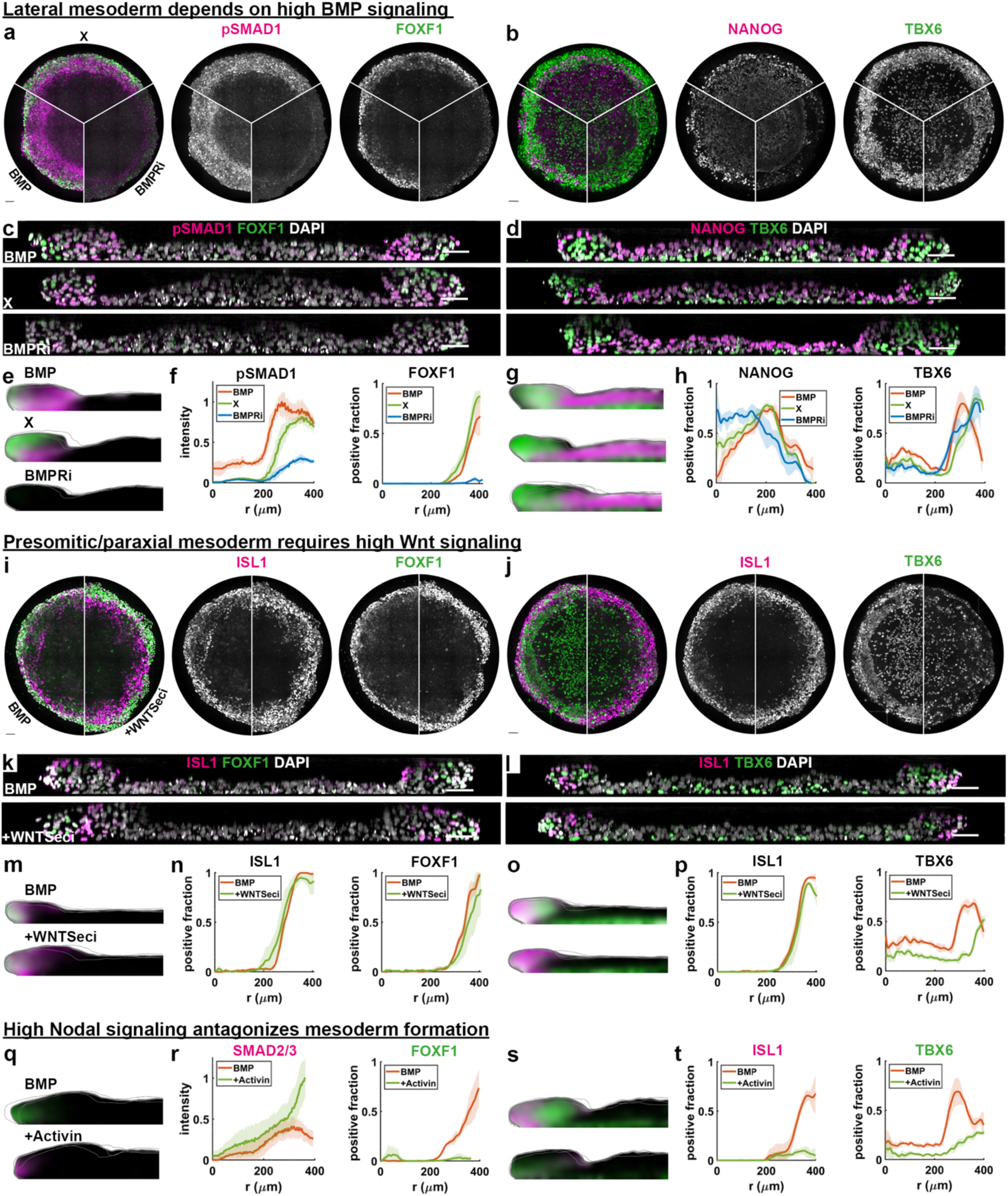
Mesodermal populations depend on distinct signaling activity. (**a-h**) Dependence of FOXF1+ (a) and TBX6+ (b) mesoderm populations as well as pSMAD1 (a) and NANOG (b) at 96h on BMP signaling after 48h. All colonies were treated with BMP4 at 50ng/ml until 48h. Then conditions were switched to BMP: BMP4 at 50ng/ml; X: only base medium mTeSR; or BMPRi: LDN193189 at 250nM. (a,b) Maximal intensity z-projections. (c,d) Cross-sections (e,g) Quantification of positive fractions in rz, and (f,h) Radial distribution of cells positive for each of the markers and levels of pSMAD1. **(i-p)** Arrangements same as (a-h). Dependence of FOXF1+ (i) and TBX6+ (j) mesoderm populations as well as ISL1 at 96h on Wnt signaling after 48h. WNTSeci: WNT secretion inhibitor (IWP2 at 5uM). **(q-t)** Arrangements same as (e-h & m-p). Dependence of FOXF1+ and TBX6+ mesoderm populations as well as SMAD2/3 and ISL1 at 96h on Activin/Nodal signaling after 48h. Scale bars 50um. Source data are provided in a Source Data file.

We next inhibited Wnt signaling after 48h. This did not affect FOXF1 expression but eliminated the TBX6 streak-like ring and significantly reduced the TBX6+ population in the center (Fig.4i,j, Supp.Fig.5f-h). We included ISL1 as a reference co-stain that partially overlaps with FOXF1 and found it was also unaffected by Wnt inhibition. Finally, we investigated Nodal signaling, as both mesodermal populations are thought to require reduced Nodal signaling, with high Nodal signaling after primitive streak differentiation generating endoderm instead^19,27,28^. We added Activin after 48h to increase Nodal signaling and observed a significant reduction in both mesodermal populations (Fig.4q-t, Supp. Fig. 6). Altogether, the results of these signaling manipulations are consistent with the proposal that our main mesoderm populations representing lateral and paraxial mesoderm.

## Discussion

We have extended culture of 2D gastruloids from 2 to 4 days and shown that during this period, development proceeds in an organized and reproducible manner that recapitulates aspects of gastrulation at later stages than have been modeled before. We observed inward directed migration from the primitive streak-like region giving rise to a paraxial mesoderm-like layer underneath the pluripotent epithelium in the center, as well as outward migration giving rise to a ring of lateral and extraembryonic mesoderm on the colony edge. These processes are self-organized and independent of exogenous BMP in the media that initiates differentiation. We showed that lineage allocation can be changed by manipulating cell signaling. By changing the balance between BMP, Wnt, and Nodal signaling, fates shift from lateral/posterior to medial/anterior. Moreover, our paraxial-like and lateral-like mesoderm closely resemble what was annotated in the CS7 human embryo as emergent and advanced mesoderm, validating these extended gastruloids as a platform that faithfully recapitulates mesoderm differentiation in the embryo.

Because this system is highly reproducible and allows tracking of cells over time, it has provided new insights into the mechanisms of gastrulation. In fact, our results suggest that the now widely used terms nascent, emergent, and advanced mesoderm^17^ may need to be reconsidered: emergent mesoderm cells that migrate inward are unlikely to give rise to advanced mesoderm cells that migrate outward later. This strongly suggests that, rather than different stages along the same lineage, these different mesoderm populations likely represent precursors of distinct lineages that emerge along a transcriptional continuum. Moreover, we showed that the advanced mesoderm likely consists of both lateral and extra-embryonic mesoderm. Altogether, we have demonstrated that extended culture of 2D gastruloids provides a new window into human mesoderm migration and patterning.

Many important questions remain. Little is known about the mechanisms driving mammalian mesoderm migration, for example whether chemo-attractants or repellents are involved, or what the mechanics are by which mesoderm moves underneath the epiblast-like layer. Lineage specification is also insufficiently understood. For example, although the FOXF1 positive mesoderm depends on BMP signaling, BMP signaling is high in a much larger region than where FOXF1 is expressed, and it is therefore unclear when and how BMP signaling causes a subset of streak-like cells to become lateral or extraembryonic versus paraxial. Another interesting question is what controls the proportions between different tissues in the system. For example, although the pluripotent layer becomes pseudo-stratified, it does not expand laterally or buckle.

To our knowledge this is the only model for human gastrulation in which different mesodermal lineages have been shown to arise in a spatiotemporally organized manner and through migration from a streak-like region. The 2D micropatterned approach is ideally suited for investigating the underlying mechanisms, as migration takes place on a micropatterned substrate that is easily manipulated, for example changing its stiffness or embedding beads for traction force microscopy^29,30^. At the same time, migration on a two-dimensional substrate is ideal for live cell imaging at resolutions required for cell tracking.

However, technical advantages and simplicity come at a price: 2D gastruloids lack many aspects of the *in vivo* gastrula. Perhaps most significant in this context is the lack of symmetry breaking to generate distinct medial-lateral and anterior-posterior axes, likely due to the absence of hypoblast which is responsible for this symmetry breaking in the embryo. That makes it difficult to interpret what it means for cells to migrate inward or outward from the streak-like ring, when comparing to migration from different anterior-posterior positions along the primitive streak. However, the primitive streak can also be converted to a ring *in vivo*, as was demonstrated for mouse and chick^31,32^. It has been argued that this represents a reversal to the ancestral model of mesoderm formation from a ring, for example as in fish, where mesoderm emerges along the margin of the embryo^33^.

The simplicity of the system and its resulting biological limitations do not only have a technical upside in making certain questions much more tractable: there are also significant advantages in following research guidelines and policies. As a non-integrated stem cell-based embryo model (SCBEM) without trophoblast or hypoblast, and without breaking rotational symmetry, the developmental potential of these structures is very limited. This ameliorates if not eliminates possible ethical concerns associated with integrated models and is compliant with federal funding policies in the U.S.^34,35^. Even in countries where funding agencies allow research with integrated models, policy guidelines being developed for SCBEM research are likely to include a requirement to use the simplest possible model that can achieve the objective. Thus, following the aphorism that all models are wrong, but some are useful^36^, as a model of the embryo 2D gastruloids are proving very useful for investigating aspects of human gastrulation that until now remained inaccessible.

## Acknowledgements

We thank Aryeh Warmflash, and Adam Helms for discussions and feedback on the manuscript, and Susana Chuva de Sousa Lopes for pointing out the connecting stalk is unaccounted for in the Tyser et al. annotation. Library prep and next-generation sequencing was carried out in the Advanced Genomics Core at the University of Michigan. This work was supported by NSF RECODE (2033654), the National Institute of General Medical Sciences (NIGMS R35GM138346), and the Branco Weiss Fellowship – Society in Science. EF was partially supported by the NIH Cellular Biotechnology Training Program (T32GM145304) and KJ was partially supported by the Michigan Pioneer Postdoctoral Fellowship and the NIH F32 Ruth L. Kirschstein Postdoctoral National Research Service Award (5F32HD108980-02).

## Data availability

Raw single cell RNA sequencing data generated in this study has been deposited in GEO (GSE262081), analyzed data will be added upon publication. Raw image data are available upon request.

## Code availability

All code for data analysis and model simulations is available on https://github.com/idse/extendedgastruloid

## Competing Interests Statement

The authors declare no competing interests.

## Methods

### Cell lines

The cell lines used were the embryonic stem cell line ESI017 (XX), and the induced pluripotent stem cell line PGP1 (XY), and a genetically modified version of the embryonic stem cell line H9 (XX) T (brachyury)-neon^37^. The pluripotency of these cells was confirmed by immunostaining of pluripotency markers OCT3/4, SOX2, NANOG. All cells were routinely tested for mycoplasma contamination, and negative results were recorded.

### Cell culture and differentiation for 2D gastruloids

Human embryonic or induced pluripotent stem cells were cultured in the commercially defined pluripotency maintaining media mTeSR1 (StemCell Technologies) on Cultrex (R&D Systems)-coated tissue culture plates. For routine cell maintenance, cell passaging was performed every 3 days. For whole-colony passaging, L7^38^ was used, while single-cell passaging was performed with Accutase. Single-cell suspensions were used to seed for all the differentiation experiments to control initial cell number.

To differentiate cells for 2D gastruloids by micropatterning, we followed the protocol in Teague et al., 2024, with an added growth period on the micropattern before BMP treatment and daily media changes. Briefly, cells were resuspended in a single-cell suspension and seeded in laminin-coated micropatterned wells in mTeSR with ROCK inhibitor (RI) (MeChemExpress, cat# HY-10583). Micropatterned wells were washed with PBS−/− 30 minutes after the initial seeding to wash off the non-attached cells. Cells were then pre-incubated for 24 hours in mTeSR before adding the BMP4 treatment by a full medium change. For all the micropatterned differentiation, full media changes with designated treatment(s) were performed every 24 hours. Experiments were performed in micropatterned 18-well Ibidi slides (cat# 81818) prepared as previously described^31^. Cell signaling reagents and doses used are listed in Supplementary Table 1.

### Immunofluorescence staining

Samples from the 18-well Ibidi slides were rinsed with PBS−/−, fixed with 4% paraformaldehyde for 30 minutes at room temperature (RT) followed by another two more times PBS−/− wash. Samples were then permeabilized with 0.1% Triton X-100 in PBS−/− for 10 minutes and blocked with blocking buffer (3% donkey serum + 0.1% Triton X-100 in PBS−/−). After blocking, samples were incubated with designated primary antibodies dilutions in blocking buffer at 4°C overnight. The next day, samples were washed with three times with PBST (0.1% Tween 20 in PBS−/−) followed by incubation with secondary antibody solution with DAPI (1ug/ml; ThermoFisher Scientific) for 30 minutes at RT without light exposure. Antibodies and their dilutions are listed in Supplementary Table 2 & 3. After staining samples were washed again with PBST two more times and stored with cover in PBS−/− with 0.01% sodium azide.

### Microscopy

Fixed sample imaging was performed with a Nikon/Yokogawa spinning dish confocal microscope with a x40 silicon oil objective using NIS Elements AR software version 5.41.02. To increase transparency, samples were treated with optical clearing buffer (FOCM) following the protocol in Zhu et al., 2019. 50% FOCM buffer was made by diluting in PBS−/− and was added immediately before imaging. Live-cell imaging was performed with an Andor Dragonfly/Leica DMI8 spinning dish confocal microscope with a x20 air objective using Andor Fusion software version 2.3.0.31 under the controlled temperature (37°C), CO_2_ concentration (5%), and humidity (>60%). The time resolution was 10 minutes. 4 z-slices were acquired with a spacing of 3.75 micron.

### Image analysis

We segmented nuclei in individual z-slices based on nuclear fluorescence (DAPI staining in fixed cells, H2B::RFP in live cells) using a pipeline we previously described^39^, which integrates two machine learning approaches: Ilastik pixel classification^40^ and Cellpose^41^ (v1). Fluorescence intensities were then calculated per nucleus as mean intensities in the 3D nuclear again with our established custom image-processing pipeline^39^. To calculate the radial fluorescence distributions in micropatterned colonies, colonies were subdivided into radial bins with equal numbers of cells, mean or median and variance were then calculated within these bins. To calculate positive fractions, markers were first thresholded based on their intensity distribution, after which the fraction of cells positive for each marker was calculated within different bins. Standard deviations were then calculated over multiple replicate colonies. To calculate the two-dimensional distribution of positive fractions in r and z, we generalized the calculation for radial profiles as described in detail in Supplementary Note 1.

### Single-cell RNA sequencing and analysis

Samples for single-cell RNA sequencing were harvested with the 10x Flex fixation kit based on the manufacturer’s manual. Single-cell RNA sequencing was performed by the University of Michigan Advanced Genomics Core. Fixed dissociated cells were counted on Luna Fx7 (Logos Biosystems). Probes were hybridized, samples were pooled and processed according to manufacturer instructions for 10X Genomics Chromium Fixed RNA Profiling Reagent Kits for Multiplexed Samples (PN 1000568). Final library quality was assessed using the LabChip GXII HT (PerkinElmer) and libraries were quantified by Qubit (ThermoFisher). Pooled libraries were subjected to paired-end sequencing according to the manufacturer’s protocol (Illumina NovaSeq XPlus). BCL Convert Software (Illumina) was used to generate de-multiplexed FASTQ files and CellRanger 7.0.1 (10X Genomics) was used to align reads and generate count matrices using the 10x supplied Human reference GRCh38 and default settings.

For comparison with the CS7 human gastrula^17^, FASTQ files were downloaded from the European Nucleotide Archive project number PRJEB40781. FASTQ files were mapped to the human reference genome GRCh38 with STAR-v2.7.11a aligner^42^ to generate count matrices. For comparison with the CS8 cynomolgus monkey embryo, (GEO dataset GSE193007) were processed with Cell Ranger 7.0.1 against the Macaca Fascicularis 5.0 genome assembly, GCF_000364345.1, which was downloaded from the NCBI website and built using Cell Ranger’s mkref function.

Further analysis was performed in Python using Scanpy^43^, Scprep^44^, and SCVI^45^. After filtering cells by library size and excluding genes with non-zero expression in less than 10 cells, we were left with 51884 cells and 16380 genes total from 6 samples (48h, 72h, 96h, in duplicate). A joint embedding of the data was generated using SCVI with cell cycle genes^46^ as nuisance variables. Differential expression analysis was performed on data that was log transformed and scaled to zero mean and unit variance using Earth Mover’s Distance. Mapping onto human and monkey embryo reference data was performed through SCVI with scANVI and label transfer from reference datasets using K-nearest-neighbors implemented in scArches. For visualization of gene expression on UMAPs in Supp.Fig.2, data were smoothened slightly with MAGIC^44^ using knn=5, t=1.

### Single cell tracking and analysis

Nuclei were segmented based on maximum intensity projections of the T-neon (brachyury) channel using a custom CellPose model^41^. Cell movements were then tracked using the label-image-detector pipeline in the Fiji Trackmate plugin^47^. Small and dim objects were filtered out, and the LAP tracker was configured with a penalty on the median T (brachyury) intensity within the nucleus. Tracks were imported into MATLAB for quantification and visualization. Relative to the colony center as the origin, the angle of deviation of a cell at a specific spatial-temporal location was determined as the signed angle between the radially inward direction and the instantaneous velocity vector. This angle was computed along the tracks for all migrating cells. The total displacement vectors map the migrating cells from their initial positions to the final positions.

### Statistics and reproducibility

All experiments were performed twice or more. All the attempts in replicating experiments yielded consistent results. Unless specifically noted, all the quantifications were performed with n=4 colonies within the same experimental condition.

### Ethics statement

We comply with all the ethical regulations in our research. We were granted approval by the Human Pluripotent Stem Cell Research Oversight (HPSCRO) Committee at the University of Michigan to work with human embryonic stem cells and human induced pluripotent stem cells.

**Supplementary Table 1:**
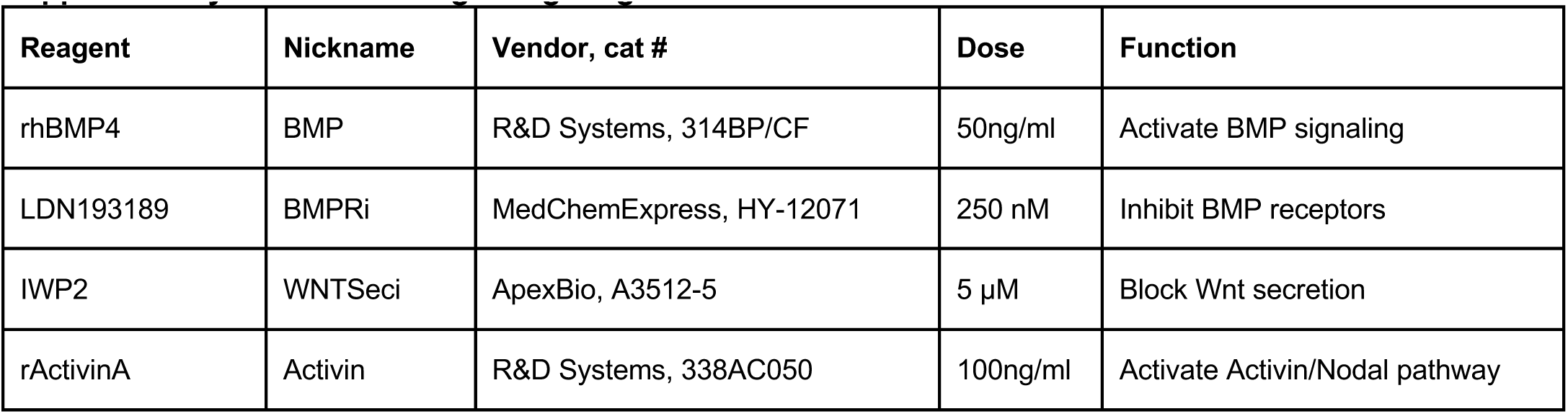
Cell signaling reagents.

**Supplementary Table 2:** Primary antibodies used for immunofluorescence.

| Antigen | Species | Dilution | Cat # | Vendor |
| --- | --- | --- | --- | --- |
| ISL1 | Mouse | 1:200 | 39.4D5 | DSHB |
| SOX2 | Rabbit | 1:200 | 3579S | Cell Signaling Tech |
| T (BRA) | Goat | 1:300 | AF2085 | R&D Systems |
| NANOG | Rabbit | 1:200 | 4903S | Cell Signaling Tech |
| NANOG | Goat | 1:100 | AF1997 | R&D Systems |
| PODXL | Mouse | 1:200 | MAB1658 | R&D Systems |
| EPCAM | Goat | 1:100 | AF960 | R&D Systems |
| E-Cadherin | Mouse | 1:200 | 610181 | BD Biosciences |
| N-Cadherin | Rabbit | 1:300 | 13116S | Cell Signaling Tech |
| EOMES | Mouse | 1:200 | MAB6166 | R&D Systems |
| MIXL1 | Rabbit | 1:150 | A17223 | ABclonal |
| TBX6 | Goat | 1:200 | AF4744 | R&D Systems |
| HAND1 | Goat | 1:200 | AF3168 | R&D Systems |
| FOXF1 | Goat | 1:300 | AF4798 | R&D Systems |
| pSMAD1/5/9 | Rabbit | 1:100 | 13820S | Cell Signaling Tech |
| TFAP2C | Mouse | 1:150 | SC-12762 | Santa Cruz Biotech |

**Supplementary Table 3:** Secondary antibodies.

| Antigen | Species | Dilution | Cat # | Vendor |
| --- | --- | --- | --- | --- |
| Alexa Fluor 488 anti-mouse | Donkey IgG | 1:500 | A21202 | ThermoFisher Scientific |
| Alexa Fluor 555 anti-rabbit | Donkey IgG | 1:500 | A31572 | ThermoFisher Scientific |
| Alexa Fluor 647 anti-goat | Donkey IgG | 1:500 | A21447 | ThermoFisher Scientific |
| Alexa Fluor 647 anti-rabbit | Donkey IgG | 1:500 | A31573 | ThermoFisher Scientific |
| Alexa Fluor 555 anti-goat | Donkey IgG | 1:500 | A21432 | ThermoFisher Scientific |

## Supplementary Figures

**Supplementary Figure 1.**
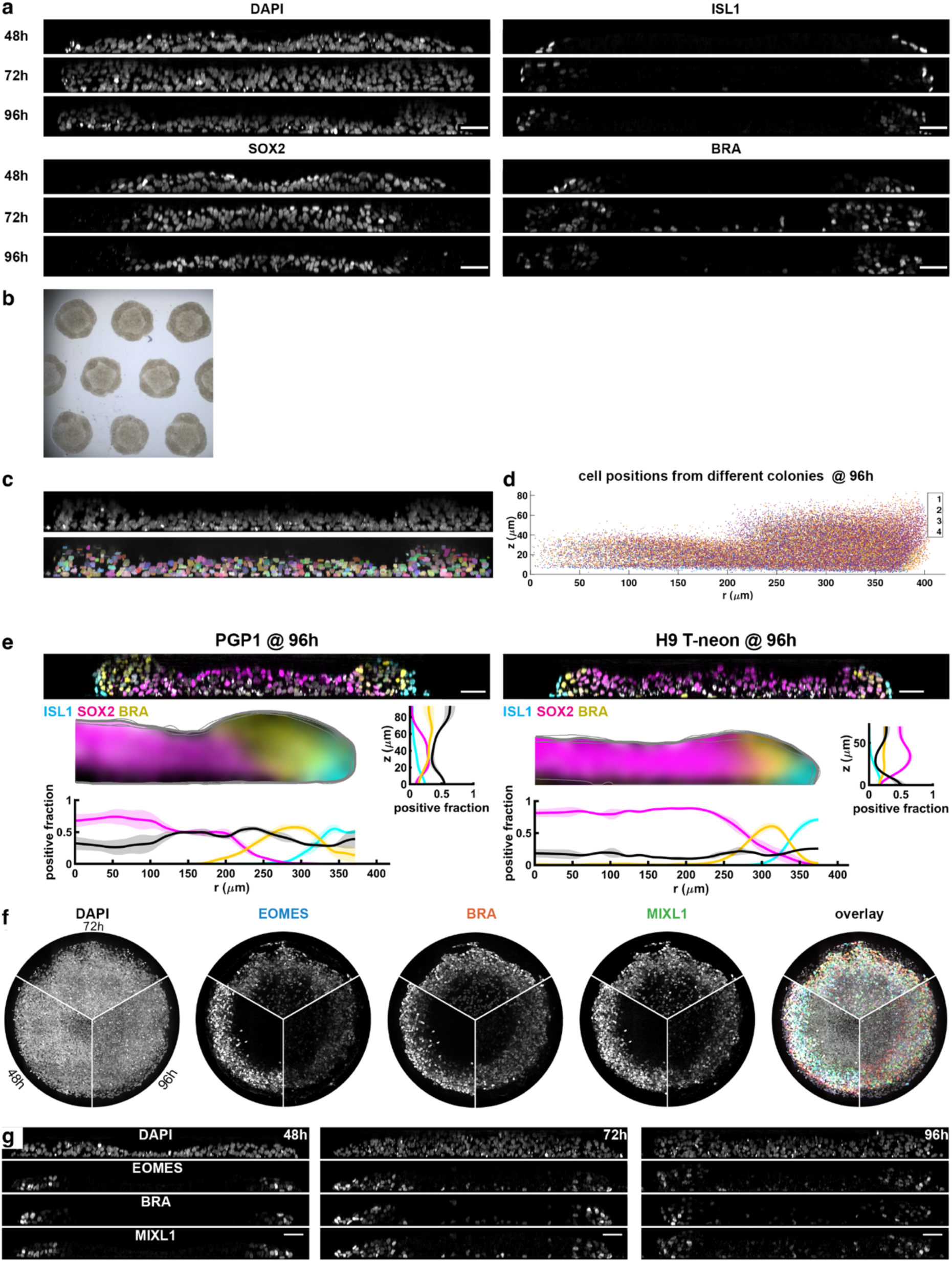
(**a**) Individual channels corresponding to Fig. 1b. **(b)** Brightfield images of 2D gastruloids at 96h. **(c)** Representative cross-section of DAPI (top) and overlay of segmentation (bottom). **(d)** Superposition of segmented cell centers from 4 replicate colonies demonstrates colony shape is quantitatively reproducible. **(e)** 96h colonies for PGP1 (XY) hiPSCs and H9 T-neon (BRA) (XX) hESCs with the same markers in Fig. 1a. **(f)** Individual channels and overlay with DAPI corresponding to Fig.1h **(g)** Individual channels corresponding to Fig.1k.

**Supplementary Figure 2.**
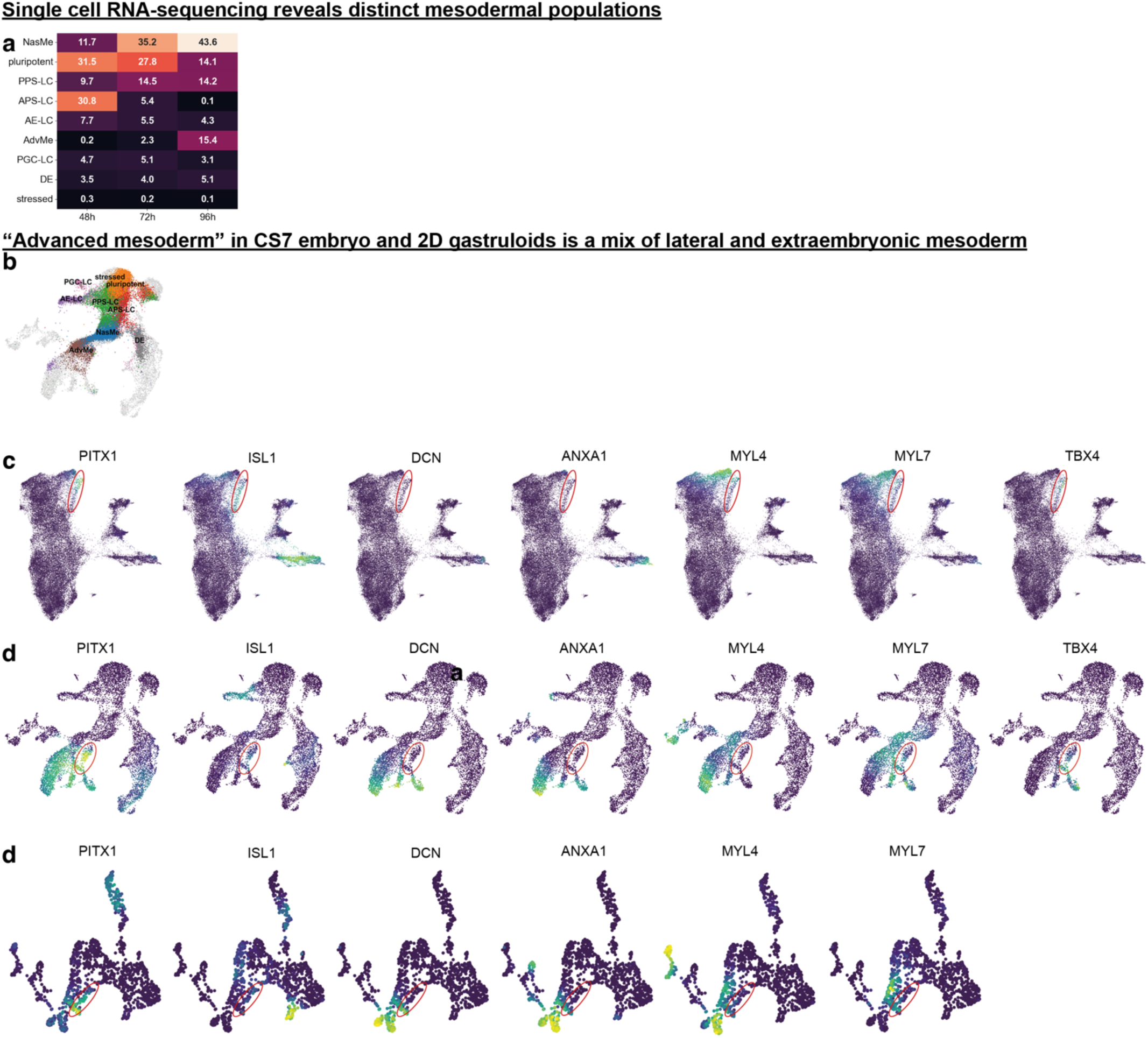
(**a**) Percentage contribution of each cluster over time. **(b)** Gastruloid clusters for complete time series mapped onto monkey embryo. **(c-e)** Expression of relevant genes in gastruloids (b), CS8 monkey (c), and CS7 human (d) shows consistent gene expression in the corresponding regions indicated by red circles. For visualization purposes, gene expression was smoothened slightly with MAGIC^44^.

**Supplementary Figure 3.**
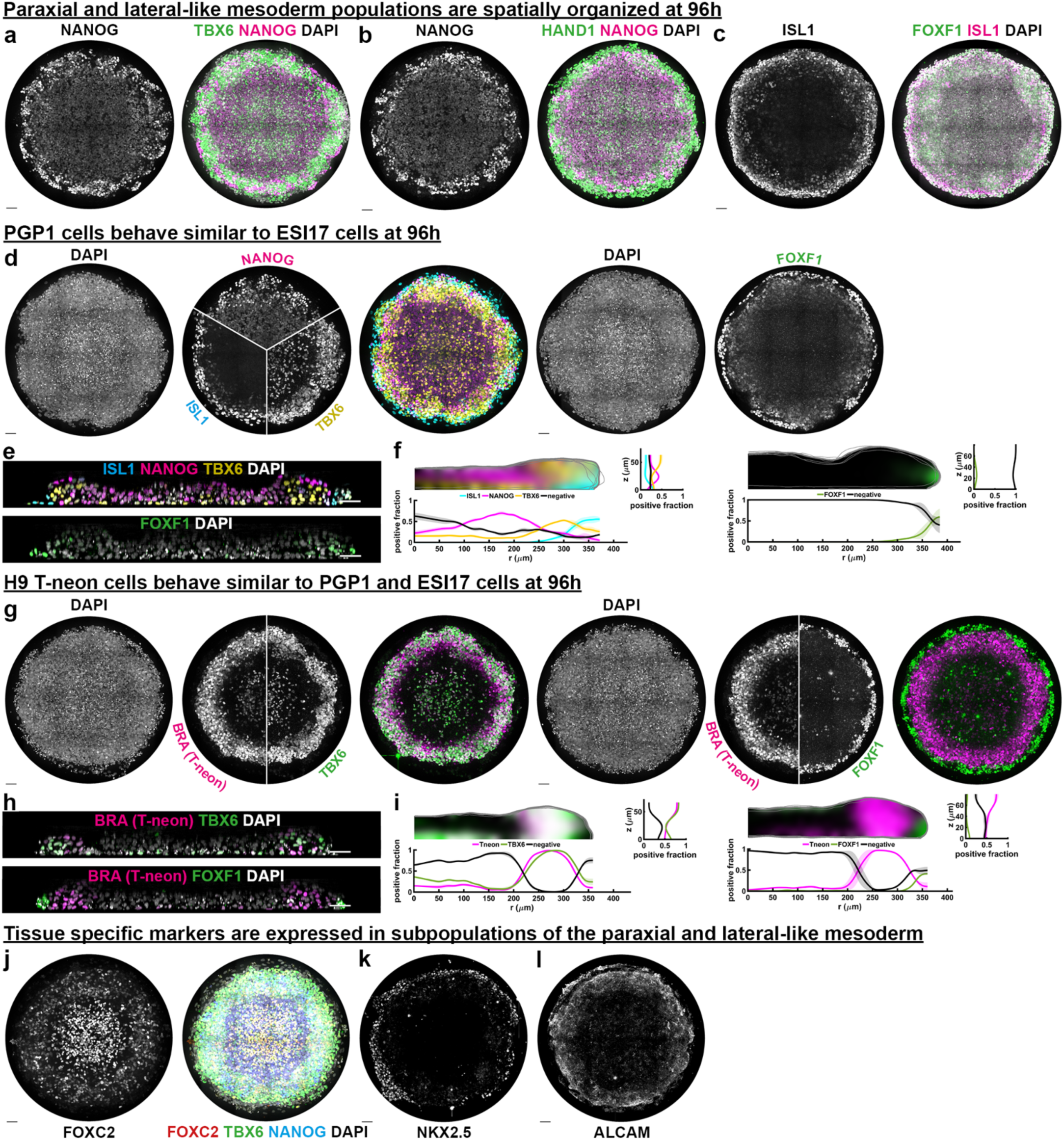
(**a-c**) Individual channels of full colonies from Fig.3a,d,g **(d)** Maximum intensity projections along z in PGP1 hiPSC line at 96h showing similar spatial patterns in marker expression as for ESI017 cells in Fig.3. **(e)** xz-cross-sections through the colony centers corresponding to (d). **(f)** Spatial distribution of positive fraction averaged over N=4 colonies for stains corresponding to (d,e). **(g-i)** Like (d-f) but for H9 T-neon hESC line at 96h. **(j-l)** individual channels corresponding to Fig. 3j,m,o, respectively. Scale bars 50um. Source data are provided in a Source Data file.

**Supplementary Figure 4.**
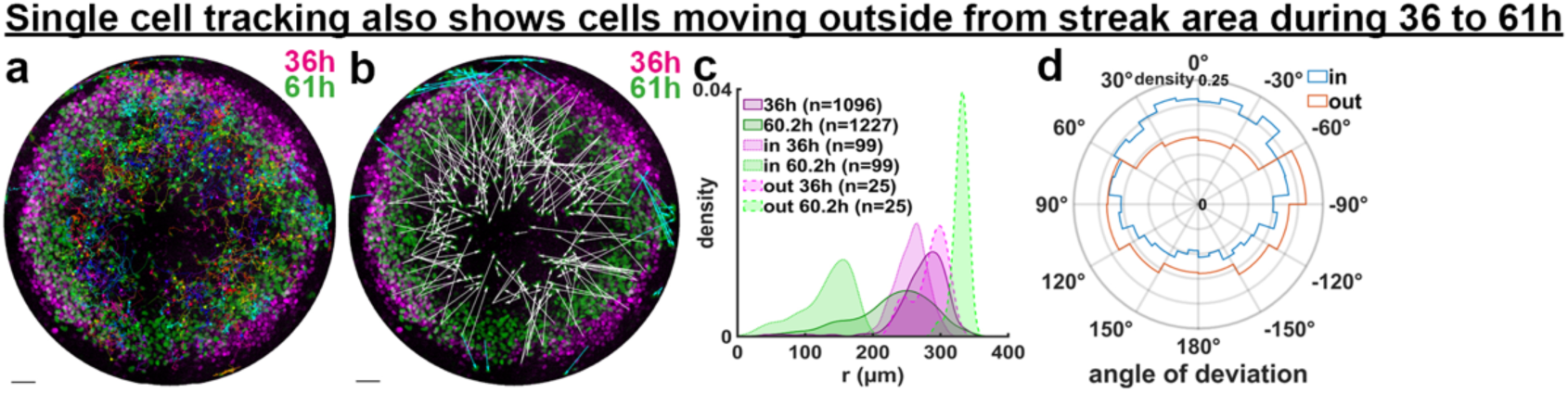
(**a**) Full tracks from 36-61h for cells that end up in the colony center overlayed on initial (magenta) and final (green) frame of T-neon (BRA) channel, corresponding displacement vectors from track start to end are shown in Fig. 3q. **(b)** Displacement vectors from 36-61h for BRA+ cells that end up in the colony center (white) or on the colony edge (cyan). **(c)** Initial (magenta) and final (green) radial distributions for BRA+ cells that end up in the center or on the edge. **(d)** Histogram of the angle of deviation from radially inward and outward motion for all track segments in (a) for cells tracked to the center (blue) or edge (red).

**Supplementary Figure 5.**
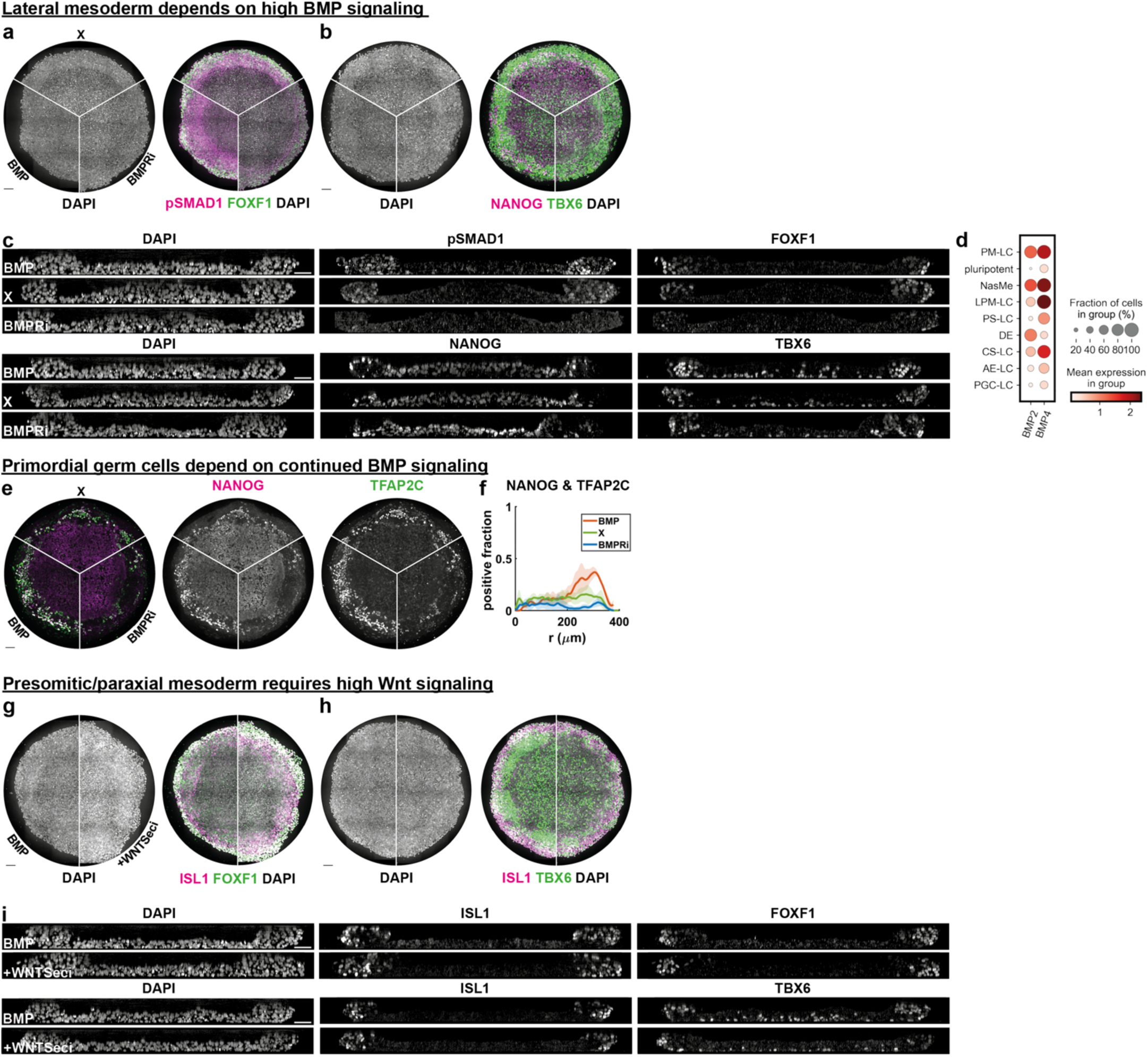
(**a,b**) DAPI and individual channels merged with DAPI images corresponding to Fig.4a,b. Conditions from 48-96h: BMP4 (BMP), mTeSR media only (X), and BMP receptor inhibitor (BMPRi) **(c)** individual channels corresponding to Fig.4c,d. **(d)** Max intensity projections along z of primordial germ cell (PGC) markers TFAP2C and NANOG under the same conditions as a-c shows loss of PGC-like cells with BMPRi after 48h. **(e)** Radial distributions of NANOG and TFAP2C co-positive fraction under different treatment conditions. **(f,g)** DAPI and individual channels merged with DAPI images corresponding to Fig.4i,j. Conditions from 48-96h: BMP4 (BMP), Wnt secretion inhibitor (WNTSeci). **(h)** individual channels corresponding to Fig.4k,l. Scale bars 50um. Source data are provided in a Source Data File.

**Supplementary Figure 6.**
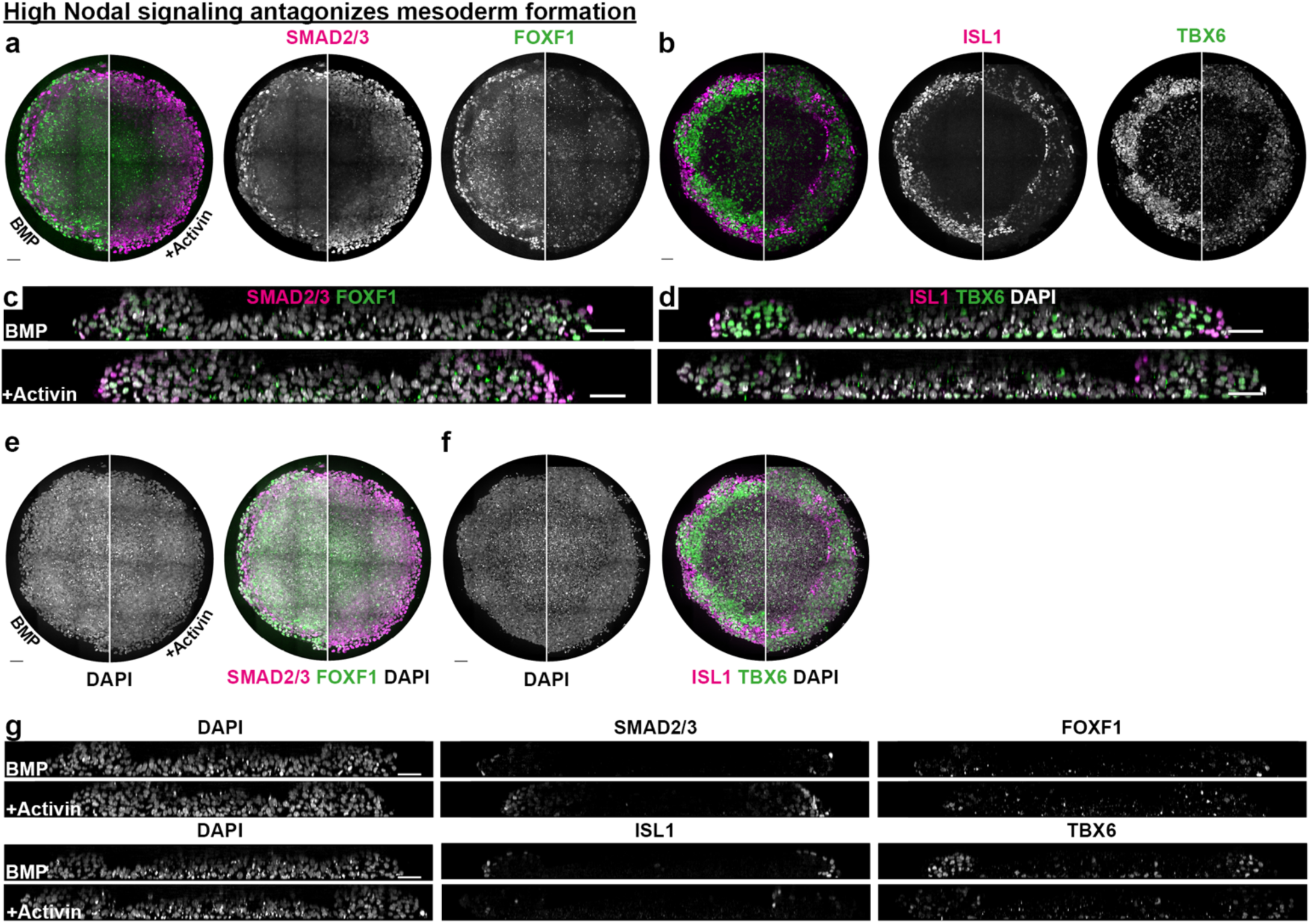
(**a-d**) Dependence of FOXF1+ (a) and TBX6+ (b) mesoderm populations as well as SMAD2/3 (a) and ISL1 (b) at 96h on Activin/Nodal signaling after 48h. All colonies were treated with BMP4 at 50ng/ml until 48h. Then conditions were switched to BMP: BMP4 at 50ng/ml; or BMP+Activin 100ng/ml. (a,b) Maximal intensity z-projections (c,d) Cross-sections. **(e,f)** DAPI and individual channels merged with DAPI images corresponding to (a,b). **(g)** Individual channels for cross-sections shown in (c,d). Maximal intensity z-projections (h) and radial distribution of endodermal cells (i). Scale bars 50um. Source data are provided in a Source Data file.

## Notes

### Competing Interest Statement

The authors have declared no competing interest.

