## Supplemental Note 1 for "Extended culture of 2D gastruloids to model human mesoderm development"

### Supplementary Note 1: Visualizing positive fractions of markers in space

February 18, 2024

We begin with histograms counts $N$ for markers $m$, using bins $r_i, z_j$ of uneven size $dr_i, dz_j$,

$$N_m(r_i, z_j), \tag{1}$$

and a total histogram $N(r_i, z_j)$. The probability of finding marker $m$ in bin $i, j$ is

$$P_m(r_i, z_j) = \frac{N_m(r_i, z_j)}{N(r_i, z_j)}. \tag{2}$$

Note that markers are not mutually exclusive, so

$$\sum_{m \in \text{markers}} P_m(r_i, z_j) := S(r_i, z_j) \geq 1. \tag{3}$$

Therefore we cannot interpret $P_m(r_i, z_j)$ as a conditional probability $P(m|r_i, z_j)$ where $m$ is a single random variable that takes on multiple discrete values. Instead we have to think of markers $m_k$ with $k$ indexing multiple random variable that each take on two values: positive or negative, so

$$\sum_{m_k \in \{+,-\}} P(m_k|r_i, z_j) = 1. \tag{4}$$

We are also interested in the marginal conditional distributions

$$P(m_k|r_i) = \frac{\sum_{z_j} N_m(r_i)}{\sum_{z_j} N(r_i, z_j)} \neq \sum_{z_j} P(m_k|r_i, z_j) \tag{5}$$

These quantities we can then interpolate on a linear grid and smooth. The smoothing part requires some care on the colony edges. If we pad with zeros where there are no cells, we will smooth all the $P(m_k|r_i, z_j)$ to zero on the edge so their sum inconsistently is less than one. To preserve the total probability, we therefore perform nearest neighbor extrapolation before smoothing, and afterwards cutt off the plot where the cell density is below 10% of its maximum.

To display these probabilities together in 2D as a heatmap on a white background, we subtract the total at each position from the maximum of that total, then add back each of the channels as an RGB vector

$$\mathrm{im} = \max(S) - S(r_i, z_j) + (P_1, P_2, P_3), \tag{6}$$

which is white when all three markers are with probability 1 or if all three are off. To resolve the ambiguity, we make areas where there are cells but all markers are off black, by calculating the fraction of cells that are negative for any marker:

$$\mathrm{im} = \mathrm{im} - P(-|r_i, r_j), \tag{7}$$

leaving the image white where all markers are on.
